# Inference of chromosome selection parameters and missegregation rate in cancer from DNA-sequencing data

**DOI:** 10.1101/2024.04.05.588351

**Authors:** Zijin Xiang, Zhihan Liu, Khanh N. Dinh

## Abstract

Aneuploidy is frequently observed in cancers and has been linked to poor patient outcome. Analysis of aneuploidy in DNA-sequencing (DNA-seq) data necessitates untangling the effects of the Copy Number Aberration (CNA) occurrence rates and the selection coefficients that act upon the resulting karyotypes. We introduce a parameter inference algorithm that takes advantage of both bulk and single-cell DNA-seq cohorts. The method is based on Approximate Bayesian Computation (ABC) and utilizes CINner, our recently introduced simulation algorithm of chromosomal instability in cancer. We examine three groups of statistics to summarize the data in the ABC routine: (A) Copy Number-based measures, (B) phylogeny tip statistics, and (C) phylogeny balance indices. Using these statistics, our method can recover both the CNA probabilities and selection parameters from ground truth data, and performs well even for data cohorts of relatively small sizes. We find that only statistics in groups A and C are well-suited for identifying CNA probabilities, and only group A carries the signals for estimating selection parameters. Moreover, the low number of CNA events at large scale compared to cell counts in single-cell samples means that statistics in group B cannot be estimated accurately using phylogeny reconstruction algorithms at the chromosome level. As data from both bulk and single-cell DNA-sequencing techniques becomes increasingly available, our inference framework promises to facilitate the analysis of distinct cancer types, differentiation between selection and neutral drift, and prediction of cancer clonal dynamics.

## 1 Introduction

Current advancements in genomics technologies have enabled researchers to examine the extent and patterns of tumor chromosomal instability. Over the past two decades, there has been great technological and computational progress in bulk DNA-sequencing (bulk DNA-seq) methods, resulting in more uniform coverage and deeper sequencing depth at a lower cost. This paved the way for large pan-cancer genomic studies, such as The Cancer Genome Atlas (TCGA) [1] and Pan-Cancer Analysis of Whole Genomes (PCAWG) [2]. The enhanced statistical power resulting from the large sample sizes has enabled identification of cancer drivers, classification of tumor subtypes, and subsequently better diagnosis and treatment decisions based on genetic biomarkers [3, 4, 5]. More recently, single-cell DNA-sequencing (scDNA-seq) technologies have emerged as a powerful method to uncover the genomic heterogeneity in individual tumors [6, 7, 8, 9]. The DNA profiles of individual cells and their inferred phylogenies also enable the analysis of how the cancers evolved over time, and which genetic features are associated with tumor expansions, metastasis and relapse [10, 9].

The application of both bulk and scDNA-seq in cancer research has led to increased understanding of the selective role of Copy Number Aberrations (CNAs) [11, 12, 13]. Defined as deletions or amplifications of large genomic regions, CNAs have been observed to enrich oncogenes and inactivate tumor suppressor genes, contributing to uncontrolled proliferation and apoptosis evasion in cancer [14]. Successful CNA detection has resulted in better patient outcome prediction [15] and personalized treatment [16, 17].

We have recently introduced CINner, an efficient algorithm for simulating chromosomal instability during tumorigenesis [18]. It allows for flexible characterization of copy numbers in individual cells, and considers both the generation and selection of diverse karyotypes. When limited to whole chromosomes, CINner extends the approach by Lynch et al. [19] to simultaneously consider missegregation rate and tissue-specific selection parameters. In this paper, we examine the problem of inferring these variables from DNA-seq data. We construct the parameter inference method based on the Approximate Bayesian Computation (ABC) framework [20], utilizing statistics that characterize the observed copy number profiles and inferred cell phylogeny. We investigate which statistics are most informative in capturing the signals of CNA heterogeneity and selection, and analyze the accuracy of the inferred parameters when applied to simulated DNA-seq data. Importantly, our study considers a combination of both bulk and scDNA-seq data, as each platform offers distinct advantages.

The large sample sizes of recent bulk DNA-seq studies enable accurate depiction of the selection landscape. However, the method offers only limited information about cancer clonality, which carries the signals for CNA rates. On the other hand, scDNA-seq captures the heterogeneity in individual tumors, but current technical and financial limits result in modest datasets that are prone to over-fitting. By combining the two data sources, we can accurately recover both missegregation rate and selection parameters. We also investigate the dependence of inference accuracy on the data sample sizes. Finally, an important open question that we assess is the fidelity of inferred statistics, and how it might impact the accuracy of our method. Specifically, the analysis of scDNA-seq data includes the inference of cell-specific copy numbers from readcount data, and deduction of a phylogeny tree that is most compatible of the inferred genomes. As the sequencing technology and computational analysis might be prone to noise, the resulting statistics might not be accurate and can impact further parameter estimation. We test the reliability of the statistics by comparing between CINner simulations and inferences from MEDICC2 [21], a phylogeny algorithm that has shown great applicability for scDNA-seq analysis.

## 2 Results

### 2.1 Inference of Copy Number Aberration rates and selection parameters in the CINner framework

We recently introduced CINner [18], an efficient algorithm to simulate the evolution of chromosomal instability (CIN) during tumorigenesis. CINner uses a birth-death process to model tumor cells [22], where a cell’s fitness and probability of division depend on its karyotype (Figure 1a). In this paper, we focus on the problem of inferring the parameters governing whole-chromosome missegregations, which have been frequently observed across different cancer types [23, 6]. The parameters of interest include *p*_*misseg*_, the probability that a missegregation event occurs in a cell division, and selection parameters {*s*_*i*_}, where *s*_*i*_ quantifies the change in cellular fitness as chromosome *i* is amplified or deleted (Figure 1b).

**Figure 1:**
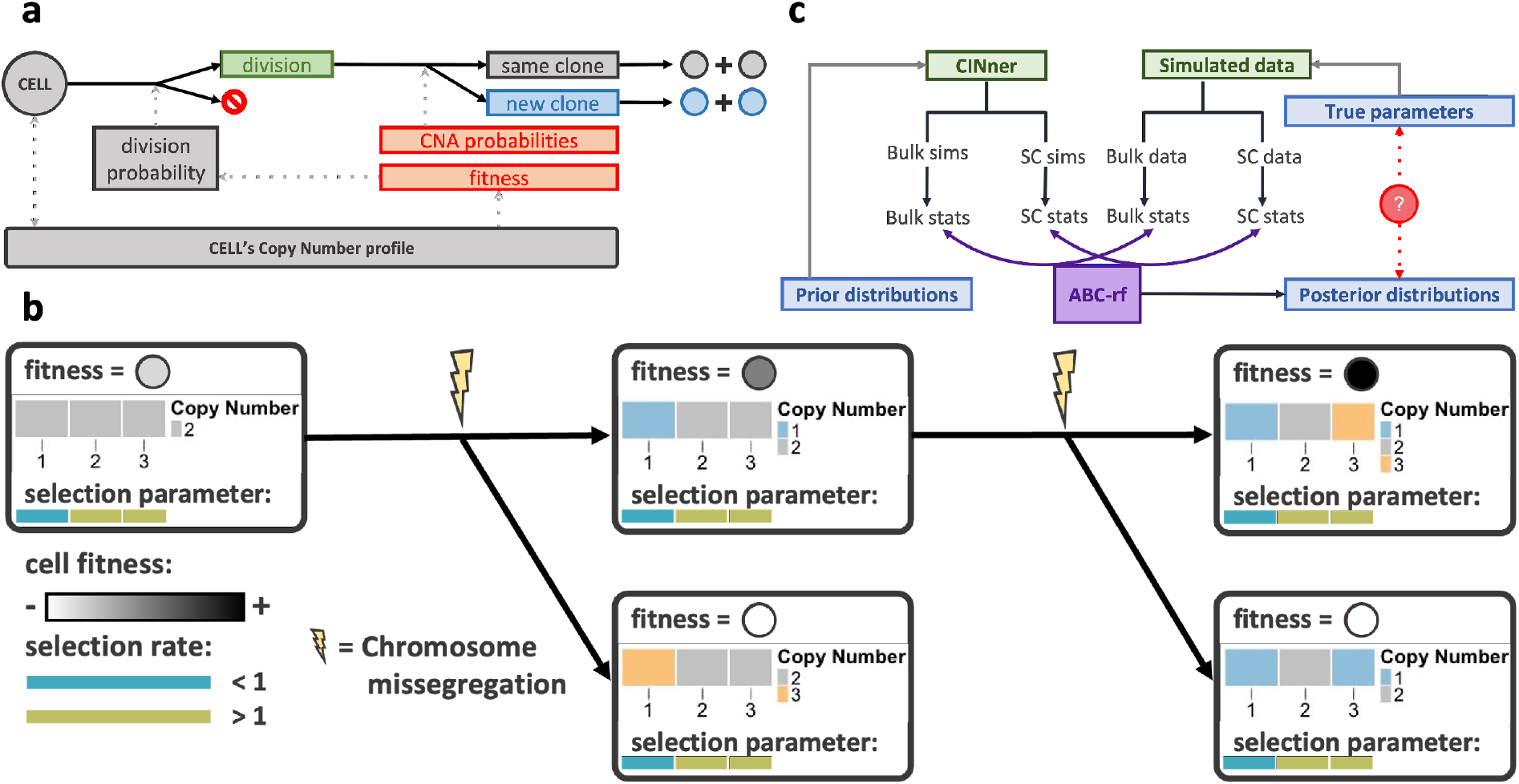
Overview of the methodology. **a**: Schematic of CINner (adapted from [18]). Cells follow a birth-death process. The probability of division depends on the cell’s fitness, determined based on its copy number profile. When cells divide, new clones may arise according to established CNA probabilities. **b**: Selection model (adapted from [18]). A cell’s fitness depends on its copy numbers and chromosome selection parameters. The fitness increases after a missegregation if the cell gains a chromosome with selection parameter > 1, or loses one with selection parameter < 1. **c**: Application of Approximate Bayesian Computation (ABC) in parameter inference. Parameters are drawn from prior distributions, then statistics are computed from bulk and scDNA-seq samples simulated with CINner. ABC-rf then determines the parameter posterior distributions that can be compared against true values.

In traditional Bayesian inference, the posterior distributions for *p*_*misseg*_ and {*s*_*i*_} are proportional to the data’s likelihood and prior probabilities [24]. However, numerical computation of the likelihood requires many simulations for each parameter set, rendering this approach too computationally expensive for our problem. Therefore, we implement Approximate Bayesian Computation (ABC), a Bayesian inference approach that replaces the likelihood by a distance function between statistics from the data and those simulated from a model. Simulation has been used to approximate the likelihood [25], and it was applied to estimate the posterior distributions of coalescence times and mutation rates from DNA-seq data in population genetics [26].

Over the last decades, ABC has been used in different fields of study to estimate parameters for complex models, especially in biology [27]. Many algorithms have been developed to improve ABC’s performance for different task requirements [28]. For our problem, we consider a wide range of statistics for DNA-seq observations, to be discussed in the next section. We utilize ABC-random forest (ABC-rf), as the algorithm is less sensitive to noise impacted by poor choices of summary statistics [29] (Figure 1c). For each parameter, ABC-rf builds a random forest from a training set consisting of sampled values from the prior distribution and corresponding simulated statistics. It then predicts the posterior distribution with regression from the random forest conditional on observed statistics from data, without requiring a metric on the statistic space.

### 2.2 Statistics for Copy Number profiles and cell phylogeny from DNA-sequencing data

The accuracy and robustness of ABC’s application in parameter inference depend heavily on the choice of summary statistics [29]. This is especially true for methods depending on a metric to compare statistics from data and model [28]. In this section, we describe several statistics for Copy Number profiles and cell phylogeny, and analyze their performance in indicating the signals of CNA rates and selection parameters. The statistics are categorized into three groups, depending on the target aspect of the data.

Several statistics estimate the heterogeneity and CNA burden from CN profiles, which we categorize as “CN statistics”. Shannon diversity index [30] measures species diversity in a given cohort based on species count and abundance, and has been widely used in genetic population studies. Given a scDNA-seq data set, individual cells can be clustered into subclones based on their CN profiles. The Shannon diversity index can then be computed from the subclone count and the sizes of each subclonal population. A more heterogeneous sample would result in higher Shannon index, and vice versa. The clonal and subclonal CNA event counts can also be calculated from CN profiles (Figure 2a). Clonal CNAs are exhibited in all cells, and therefore assumed to have occurred before the Most Recent Common Ancestor (MRCA) in the cell phylogeny. Subclonal CNAs occur after the MRCA and are carried only by a subgroup of cells. We define the subclonal CNA count as 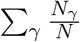, in which each CNA *γ* is weighted by *N*_*γ*_, the number of cells carrying it, against the total cell count *N*. Finally, we directly compute the distances between CINner simulations and DNA-seq data cohorts. For a bulk DNA-seq set containing *n* samples, we produce *n* simulations. The distance matrix *A* ∈ R^*n*×*n*^ indicates the precision of the simulation cohort, where *A*(*i, j*) is the Euclidean distance between CN profiles of simulation *i* and DNA-seq sample *j*. The optimal transport algorithm [31, 32] then calculates the minimal distance from the simulation cohort to the data cohort, which we refer to as the bulk DNA distance (Figure 2b). A similar strategy is used to compute the scDNA distance (Figure 2c). Each data sample *j* can be represented as 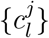, where 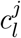 is the CN profile of cell *l*. Similarly, the CN profiles from simulation *i* are defined as 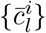. We define the distance matrix 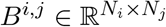, where *B*^*i,j*^(*l*_1_, *l*_2_) is the Euclidean distance between 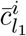 and 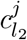. The entry *A*(*i, j*) in the distance matrix *A* is then the optimal transport distance measured from *B*^*i,j*^, and a final application of optimal transport on *A* produces the scDNA-seq distance. This approach can be computationally prohibitive, as recent scDNA-seq cohorts contain up to several thousand cells. To decrease the runtime, we define *B*^*i,j*^ for each pair of subclones in simulation *i* and data sample *j*. The value for *A*(*i, j*) then results from the optimal transport where the probability distributions for simulation *i* and data sample *j* are weighted for the subclonal cell counts.

**Figure 2:**
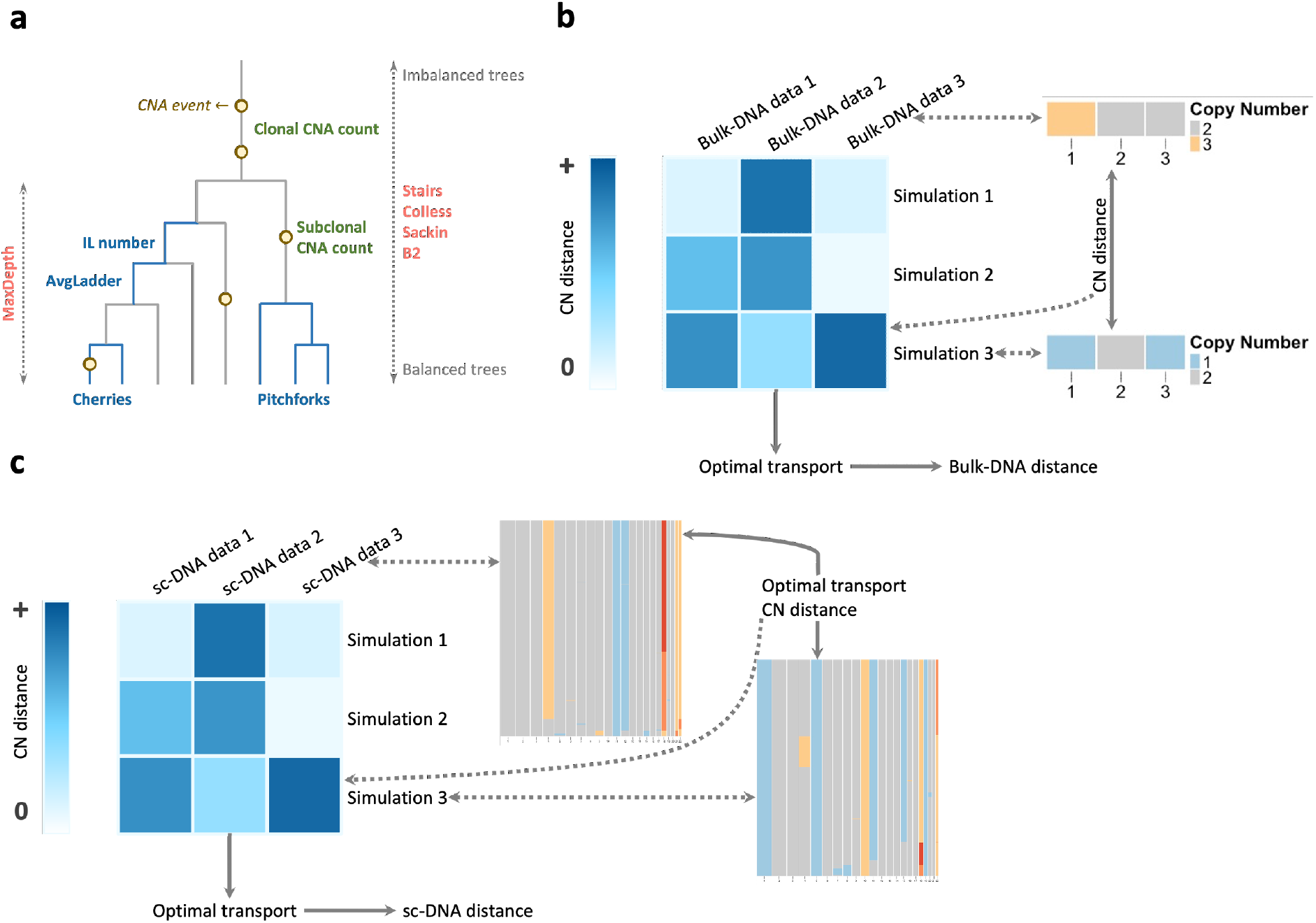
Some statistics quantifying the similarity between DNA-seq data and simulations. **a**: Statistics from CN data and cell phylogeny are grouped into CN statistics (green), phylogeny tip statistics (blue) and phylogeny balance statistics (red), depicted on a representative phylogeny tree. **b**: Bulk DNA distance is defined as the optimal transport cost from simulations to data samples. The distance matrix consists of Euclidean distances between each pair of samples. **c**: The scDNA distance is similarly based on the optimal transport from simulations to data samples. The distance between each pair of samples requires first finding the optimal transport between the sampled cells.

We divide the cell phylogeny statistics into two groups, the “tip statistics” and the “balance statistics” (Figure 2a). The tip statistics are associated with the leaves in the phylogeny tree. A cherry is defined as two leaves that merge directly with each other, and a pitchfork is a group of three leaves merging into one internal node. Cherry and pitchfork counts are the normalized numbers of these structures in the entire tree [33]. A ladder is defined as a sequence of internal nodes where each node has exactly one direct descendant. IL number and average ladder are the count of ladders and the average ladder length in the phylogeny tree, respectively [34]. In contrast to the tip statistics, the balance statistics quantify whether the tree is balanced or imbalanced. Max depth is the height of the phylogeny tree when branch lengths are normalized, which is smaller for a more balanced tree [35]. Stairs measures the proportion of subtrees that are imbalanced [34]. Colless is the sum of balance values among all internal nodes, where the value for each internal node is the absolute difference between sizes of clades stemming from it [34]. Both Sackin and Colless indices measure the imbalance extent of trees. In contrast, B2 is a balance index. B2 for tree *N* is defined as *B*_2_(*N*) = − Σ _*𝓁*∈*L*_ *p*_*𝓁*_ log_2_ *p*_*𝓁*_, where *p*_*𝓁*_ is the probability of ending at leaf *𝓁* conditioned on traveling from the root with equal probabilities at each merge [36].

In order to evaluate the effectiveness of each statistic in capturing the signals of CNA rates and selection parameters, we compute the statistics from CINner simulations with varying parameters. The chromosome selection parameters for each simulation are sampled from Uniform(1*/*1.2, 1.2), and the missegregation probability is sampled as log_10_(*p*_*misseg*_) ∼ Uniform(−5, −3). For each parameter set, we simulate 100 bulk samples and 50 scDNA samples, then compute the statistics as described above. Afterward, we compute the correlation between each parameter and the mean and variance of each statistic from 100,000 simulations. If a statistic strongly correlates with a parameter, it is potentially a good candidate for ABC parameter inference. We note that, as the selection parameters only affect the chromosomes that they are assigned to, they have minor impact on genome-wide statistics. Therefore, the CN statistics (e.g. Shannon diversity index, clonal and subclonal CNA counts, and CN distances) are computed based only on chromosome *i* when compared agaisnt selection parameter *s*_*i*_. In contrast, CNA rates affect all chromosomes, therefore they are compared against CN statistics computed for the entire genome. The other statistics are based on cell phylogeny, which cannot be segregated for individual chromosomes. Therefore, the same values are compared with both selection parameters and missegregation probability. Finally, the CN distances require direct comparison to the data samples. Therefore, we simulate 100 bulk samples and 50 scDNA samples to serve as the DNA data, with ground-truth parameters *p*_*misseg*_ = 2 × 10^−4^ and *s*_*i*_ ∼ Uniform(1*/*1.15, 1.15). Because the CN distances measure the proximity between entire cohorts of data samples and simulations, they can be used at once without other summary statistics such as mean or variance.

The correlations between the DNA-seq statistics and CINner parameters are presented in Figure 3. All of the CN statistics are strong signals for the missegregation probability. As the probability increases, the samples become more heterogeneous and contain more aneuploidies both at clonal and subclonal levels, Therefore, the Shannon index and missegregation counts are positively correlated with the missegregation probability. Compared to the CN statistics, the correlations between the phylogeny tip statistics and the missegregation probability are extremely weak. This indicates that these statistics are more representative of the sample size than of the heterogeneity associated with the scale of CNA rates. Specifically, if the subclone count is small compared to the sample size, then the cherry count in the phylogeny tree is approximately the sum of cherry counts in distinct subclones. Because the cells in each subclone are identical, the count only depends on the subclonal population size. The same is true for pitchforks and ladders, rendering these statistics ineffective of capturing the heterogeneity and selection from genomic data. Finally, the mean phylogeny balance indices correlate strongly with the missegregation probability. As the missegregation probability is elevated, there are increasingly more subclones with distinct fitness competing to expand, resulting in higher imbalance in the cell phylogeny. Therefore, the correlation is negative if the index increases when the trees are more balanced (B2) and positive if the index rises for more imbalanced phylogony (Colless and Sackin indices, stairs, and max depth). Moreover, the variance of B2 also correlates with the missegregation probability.

**Figure 3:**
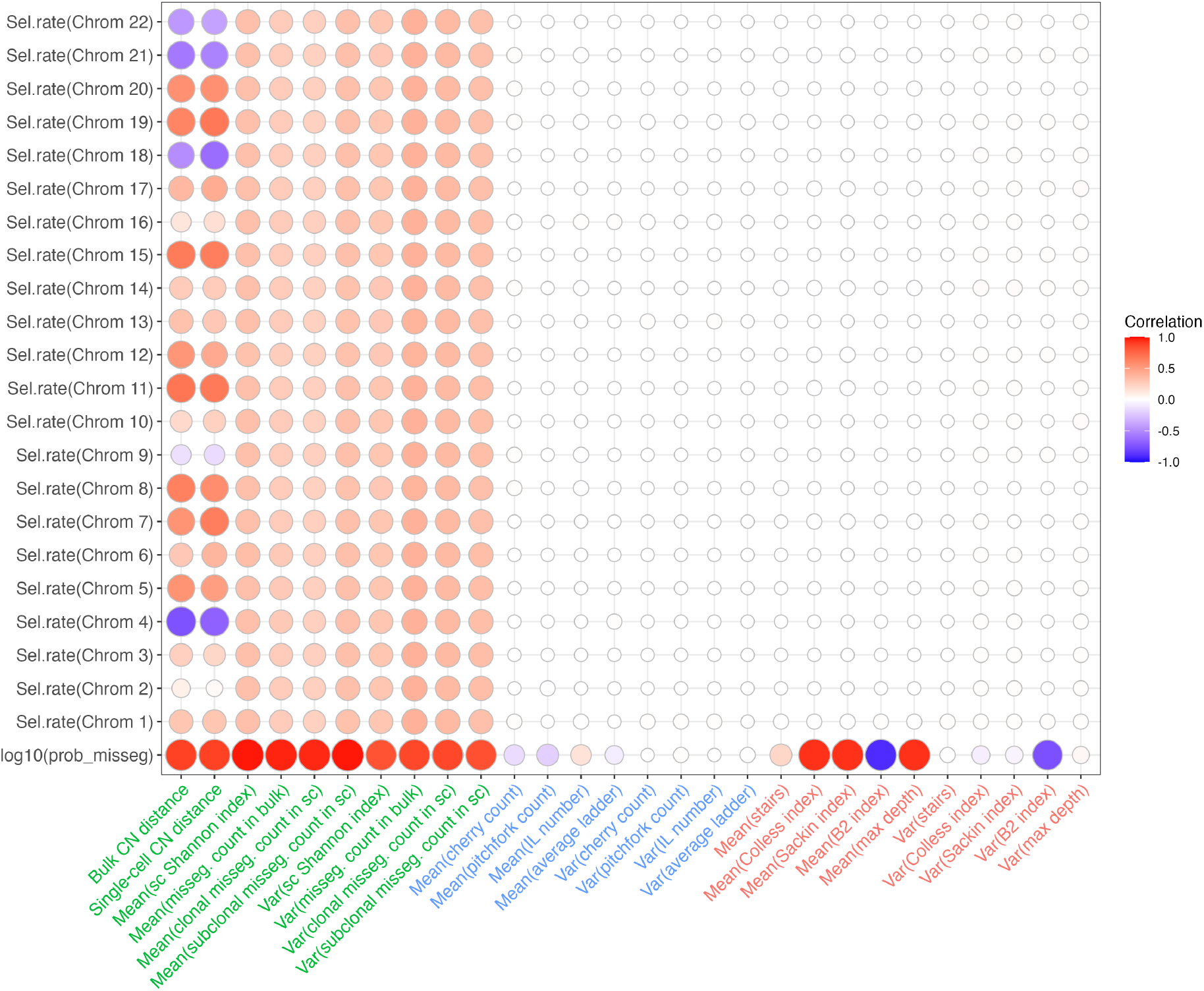
Correlations between sample statistics and model parameters. Statistics are categorized into CN statistics (green), phylogeny tip statistics (blue) and phylogeny balance statistics (red). Size and color of each circle indicate the correlation.

In contrast, only CN statistics strongly correlate with the chromosome selection parameters. Nevertheless, the correlations are weaker than between CN statistics and missegregation probability. This is because the aneuploidy level in each chromosome is significantly lower than in the entire genome. Therefore, the CN statistics for specific chromosomes are sparser than the genome-wide equivalences. As selection parameters increase, subclonal competition is intensified, and only the cells harboring the most favorable karyotypes can expand, resulting in higher missegregation counts both clonally and subclonally. The correlation signs of the CN distances depend on the individual chromosomes. If a chromosome *i* has ground-truth selection parameter *s*_*i*_ ≫ 1 (e.g., chromosomes 4, 18, 21 and 22, Figure 4a), it is frequently amplified in the observed DNA samples. As the selection parameter increases in CINner, the simulated samples also regularly exhibit gains of this chromosome, reducing the CN distances, therefore the correlation is negative. On the other hand, if the ground-truth selection parameter *s*_*i*_ ≪ 1 (e.g. chromosomes 11, 15 and 19), the observed DNA samples frequently display lower copy numbers for chromosome *i*. If the CINner simulations have higher selection parameters, the increased gain counts of chromosome *i* expand the CN distances and lead to positive correlations. The phylogeny-based statistics, including the tip statistics and even the phylogeny balance indices, have no correlation with individual selection parameters. This is because the selection parameter of one chromosome has minimal impact on the entire cell phylogeny tree. As a result, we expect that the application of these statistics is inefficient in uncovering the selection landscape from DNA data.

**Figure 4:**
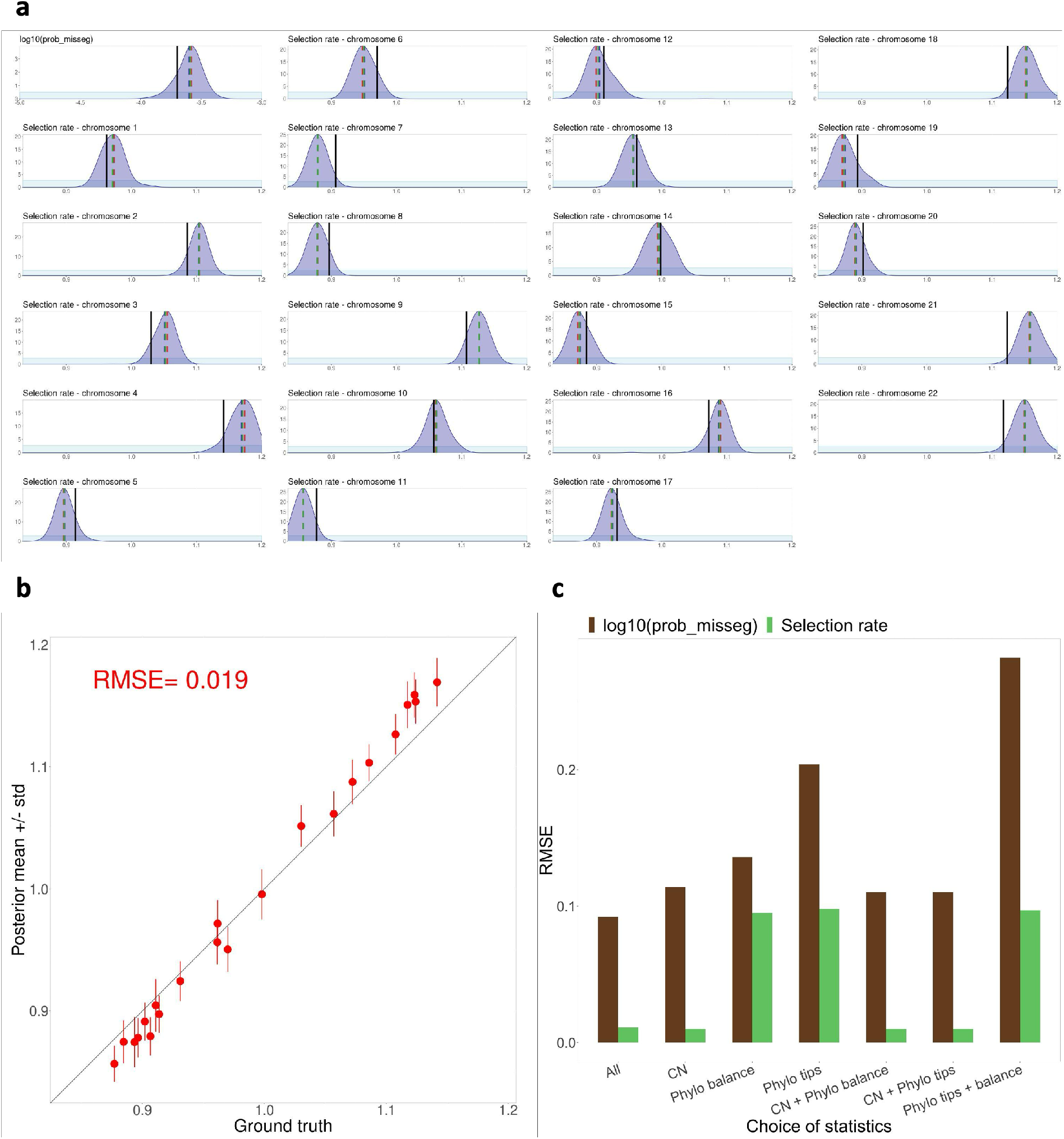
Parameter inference results with ABC-rf. **a**: Prior (light blue) and ABC-rf posterior (dark blue) distributions for each parameter, with posterior mean (blue line), mode (red line), median (green line) compared against ground truth value (black line). Inference for *p*_*misseg*_ utilizes all genome-wide statistics, inference for selection parameters uses only chromosome-specific CN statistics. Simulated data = 100 bulk and 50 scDNA-seq samples, ABC-rf training set from 100,000 CINner simulations. **b**: Correlation between ground truth and posterior mean values for all chromosome selection parameters (+/-standard deviation). Root mean square error (RMSE) computed for all selection parameters. **c**: RMSE of the posterior mean values, depending on statistic groups used in inference.

### 2.3 ABC-based parameter inference accurately recovers selection and CNA rates

We construct an ABC-based parameter inference method to recover the missegregation rate and selection parameters from a mixture of bulk and scDNA-seq data cohorts. To test the algorithm, we use CINner to create a simulated dataset by combining *N*_*bulk*_ = 100 bulk and *N*_*sc*_ = 50 scDNA samples, using ground truth parameters as described previously (*p*_*misseg*_ = 2× 10^−4^ and *s*_*i*_ ∼ Uniform(1*/*1.15, 1.15)). A training set is built from 100,000 CINner simulations, with prior distributions log_10_(*p*_*misseg*_) ∼ Uniform(−5, −3) and *s*_*i*_ ∼ Uniform(1*/*1.2, 1.2). Each CINner simulation consists of creating *N*_*bulk*_ + *N*_*sc*_ independent samples with the selected parameters, then computing the statistics as described above. Finally, we train ABC-rf [29] on the library and use the random forest to infer the posterior distribution for each parameter. To evaluate the quality of the inference, we compute the root mean square error (RMSE) [37]. Let {*α*_1_, …, *α*_*n*_} and {*β*_1_, …, *β*_*n*_} be the ground truth and inferred values for *n* parameters. The RMSE is then defined as

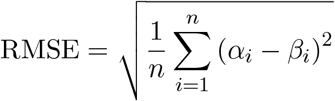

As described in the previous section, the genome-wide impact of the CNA rates means that their signals are contained in all CN and phylogeny-based statistics (with the exception of statistics based on scDNA-seq phylogeny tips). In contrast, the local effect of selection parameters renders only CN-based statistics appropriate for inference (Figure 3). Therefore, our inference for *p*_*misseg*_ employs all statistics, and the inference for each *s*_*i*_ uses only the CN statistics. The posterior distributions (Figure 4a) of either log_10_(*p*_*misseg*_) or each *s*_*i*_ exhibit almost identical mode, mean and median. Moreover, the posterior peaks are centered close to the ground truth values. Specifically, the mode, mean, and median of log_10_(*p*_*misseg*_) are −3.58, −3.6, −3.59, which are slightly higher but close to the ground truth value of log_10_(2 × 10^−4^) ≈ −3.69. Similarly, the inferred *s*_*i*_’s are close to the ground truth, with the RMSE of posterior means being 0.019 (Figure 4b). The inference tends to be slightly more extreme than the true values: if true *s*_*i*_ > 1, then inferred 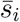 is often greater than *s*_*i*_, and vice versa. Combined, these results indicate that our inference method, based on ABC and employing appropriate statistics for each parameter class, can recover the missegregation rate and the selection landscape driving cancer evolution from DNA-seq data.

We further examine the performance of our inference method when the statistics are chosen differently (Figure 4c). As expected, for fitting *p*_*misseg*_, ABC-rf using only CN statistics performs the best, followed by phylogeny balance statistics and much higher RMSE when using only phylogeny tips measures. Intriguingly, the inference with the combination of phylogeny tips and balance statistics fares even worse than ABC-rf using either group individually. One possible explanation is that there are too few signals contained in the balance statistics to offset the increased noise introduced by the tip statistics. Note that ABC-rf is relatively insensitive to noisy statistics, compared to other ABC-based methods [29]. Algorithms such as sequential Monte Carlo (SMC) [38] or Markov Chain Monte Carlo (MCMC) [39], which use a metric to compare statistics between the model and data, would likely suffer more from the increased noise level of these measures. Finally, our chosen statistics set, consisting of all three groups, results in the lowest RMSE in inferred *p*_*misseg*_.

Similarly, the inferred selection parameters incur the lowest RMSE when CN statistics are employed, either unaccompanied or combined with phylogeny-based statistics. Because the latter is essentially noise, the inference using all statistics is similar to only using CN statistics. ABC-rf’s analysis of variable importance (Figures S1, S2 and S3) confirms that the CN statistics and the mean phylogeny balance indices are most important in inferring *p*_*misseg*_ and *s*_*i*_’s.

### 2.4 Sensitivity analysis

We showed that our inference method can recover the missegregation probability and selection parameters from a cohort of 100 bulk DNA-seq and 50 scDNA-seq samples. However, due to many constraints, the available bulk DNA-seq samples for specific cancer types can be of much smaller sizes [1]. Additionally, despite recent advances, single-cell DNA sequencing remains expensive and technologically challenging, and existing data cohorts rarely consist of more than a few samples [6, 9, 10]. Therefore, we examine the impact of the sample sizes on the accuracy of the inferred selection landscape. For a given {*N*_*bulk*_, *N*_*sc*_}, we use CINner to simulate *N*_*bulk*_ bulk DNA-seq samples and *N*_*sc*_ scDNA-seq samples, using similar ground truth parameters as in previous sections. We then infer *p*_*misseg*_ and selection parameters *s*_*i*_ from these samples, and compute the RMSE of the inferred *s*_*i*_’s, as well as the standard deviation in their posterior distributions. Lower RMSE indicates that the inferred selection parameters are accurate, and lower standard deviation implies that there is less uncertainty in the inference. We first examine the inference accuracy as dependent on the scDNA-seq cohort size. We fix *N*_*bulk*_ = 100 and vary *N*_*sc*_ = 5, 10, …, 50. Unsurprisingly, both RMSE and standard deviation decrease as *N*_*sc*_ increases (Figure 5a). This is likely because many of the statistics used in our method are based on CN profiles and phylogenies from the scDNA cohort. We note that although having very low *N*_*sc*_ incurs higher standard deviation in the selection parameters, the RMSE is still acceptable. This is probably because the signals from bulk samples (given that there are a sufficient number of them) can compensate for the low scDNA count in capturing the selection landscape, albeit with higher uncertainty. The standard deviation decreases as the scDNA sample size increases and plateaus at *N*_*sc*_ ≈ 20, and the RMSE continues decreasing up to *N*_*sc*_ ≈ 40.

**Figure 5:**
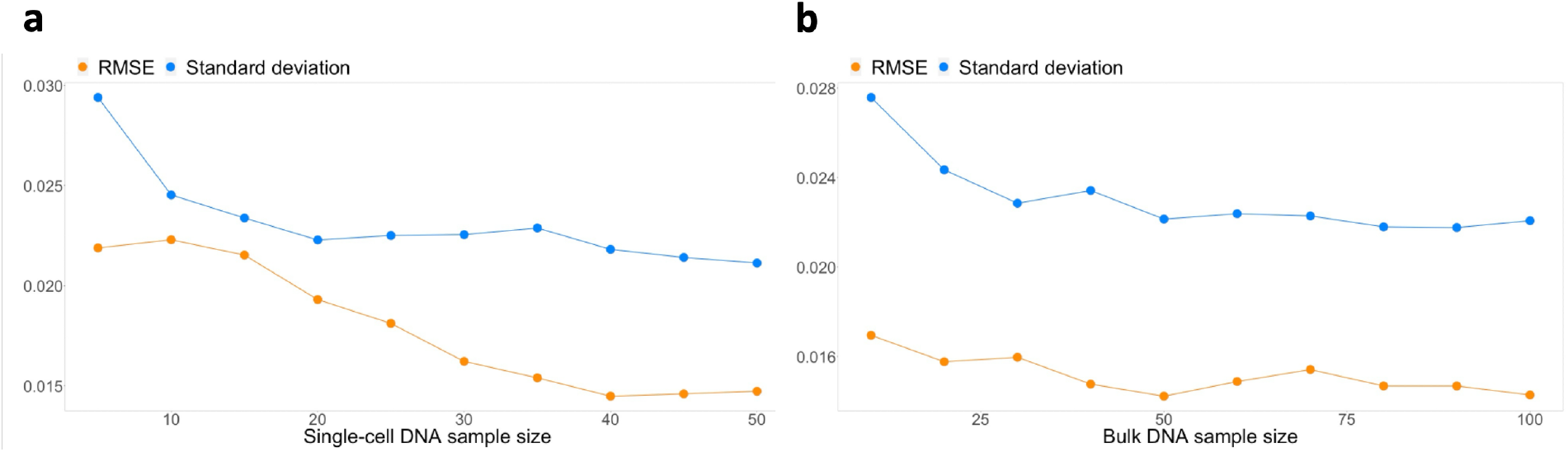
Dependence of inference accuracy on sample sizes. RMSE and standard deviation of selection parameter posterior means, as a function of scDNA-seq (**a**) and bulk DNA-seq (**b**) sample sizes.

Next, we fix *N*_*sc*_ = 50 and vary *N*_*bulk*_ = 10, 20, …, 100. The selection parameter standard deviation decreases up to *N*_*bulk*_ ≈ 30, but the RMSE only decreases slightly (Figure 5b). That the inference method depends less on *N*_*bulk*_ than on *N*_*sc*_ could be because there are only a few statistics in our method that are based on bulk DNA-seq data, namely the total missegregation counts and the distance between CN profiles in the data cohort and the CINner samples (Figure 2b). Another possible interpretation is that for adequately large *N*_*bulk*_, the bulk DNA samples can already capture the chromosome-specific CNA signals required to construct the CN distance. As a result of either or both explanations, the RMSE and standard deviation stabilize for *N*_*bulk*_ ≥ 50.

Overall, the performance of our inference method in uncovering the selection landscape reaches its peak for data cohorts consisting of as low as 50 bulk DNA-seq and 40 scDNA-seq samples. There already exists data of these scales for certain cancer types [2, 40]. More importantly, the performance is only moderately worse for much smaller data cohorts. As both sequencing technologies become more widely available, our method and its insights could become instrumental in understanding the selective forces and CNA mechanisms that drive tumorigenesis.

### 2.5 Impact of phylogeny inference accuracy on single-cell DNA statistics

The scDNA-seq statistics employed in our method are based on CN profiles and cell phylogeny inferred from sequencing readcount data. The phylogeny inference is particularly challenging, as the observed CN profiles can typically be explained by different evolution models [41]. In this section, we investigate the impact of phylogeny inference error on scDNA-seq summary statistics.

We create 1,000 CINner simulations, each with log_10_ (*p*_*misseg*_) sampled from Uniform (−5, −3) and chromosome selection parameters from Uniform(1*/*1.15, 1.15). We then apply MEDICC2 on the simulated single-cell CN profiles (Figure 6a). MEDICC2 reconstructs the phylogeny and infers the ancestral genomes from somatic CNAs, by computing the minimum-event distance between each pair of cells using a weighted finite-state transducer framework [21]. MEDICC2 produces a phylogeny tree rooted in the diploid genome, with CNAs occurring on specific branches such that the tips recover the observed CN profiles in the sample.

**Figure 6:**
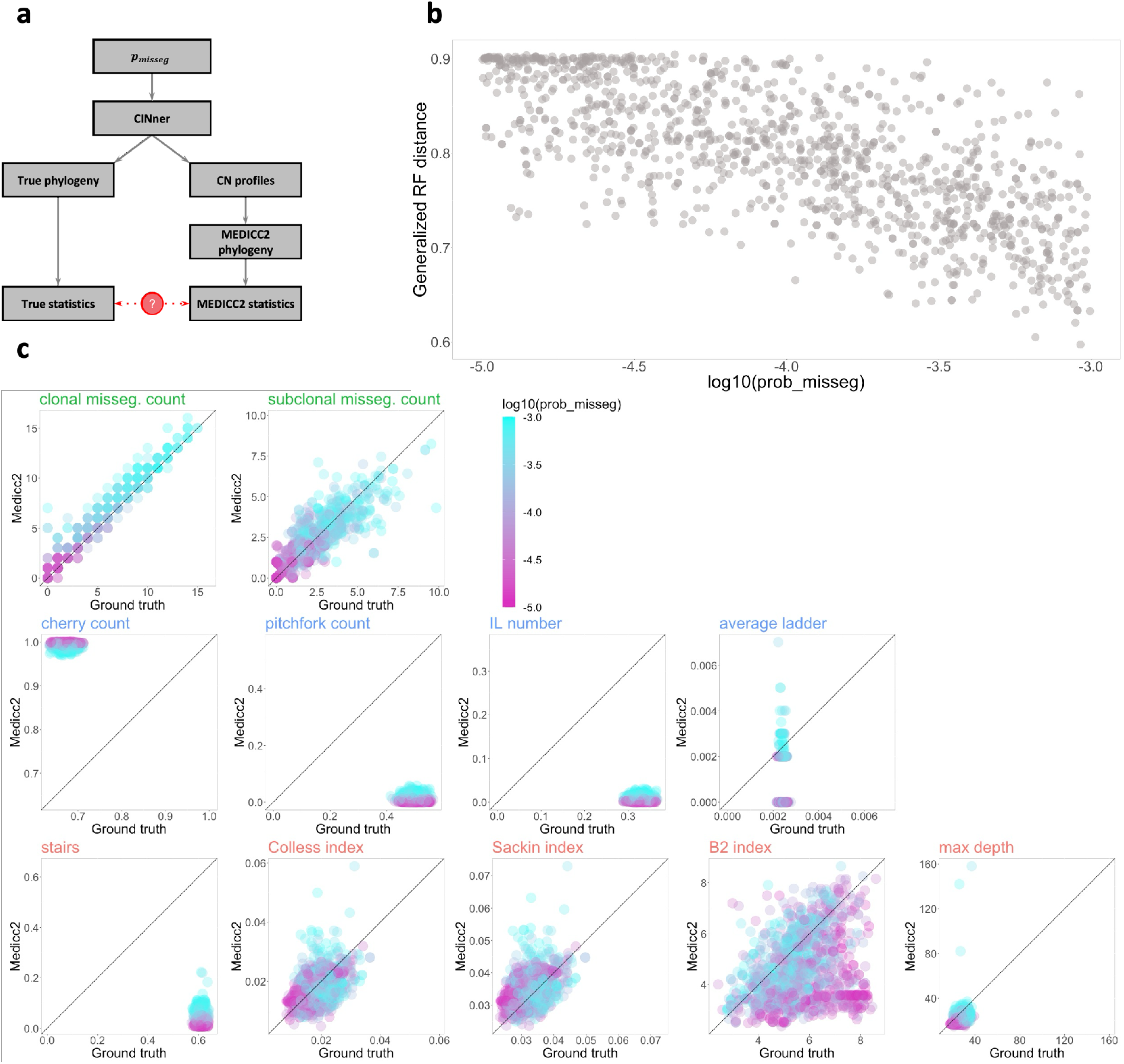
Accuracy of phylogeny statistics inferred from MEDICC2. **a**: Study schematic. Statistics computed from the true phylogeny from each CINner simulation are compared against statistics estimated from the MEDICC2-inferred tree. 1,000 simulations are performed for this study, each with log_10_ (*p*_*misseg*_) ∼ Uniform (−5, −3). **b**: Generalized Robinson-Foulds distances between true and MEDICC2 phylogeny, against corresponding *p*_*misseg*_. **c**: Comparison between statistics computed from true and MEDICC2 phylogeny. Color of each point denotes *p*_*misseg*_.

We first compare the true phylogeny against the tree inferred by MEDICC2 by using the generalized Robinson-Foulds distance [42] (Figure 6b). Unsurprisingly, the MEDICC2 inferred tree becomes more accurate as *p*_*misseg*_ increases. This is because MEDICC2 cannot stratify a group of cells if they have the same CN profiles. As *p*_*misseg*_ increases, there are more CNAs segregating distinct cells, and the MEDICC2 phylogeny becomes more resolved and closer to the ground truth.

We then analyze the summary statistics computed from MEDICC2 phylogeny (Figure 6c). The counts of clonal and subclonal missegregations, which require assigning each event on the phylogeny tree, are largely in agreement with the true values. As expected, increasing *p*_*misseg*_ results in higher missegregation counts. However, the accuracy of MEDICC2 inferences does not depend on the value of *p*_*misseg*_. This suggests that these statistics can reliably indicate the level of aneuploidy observed in the sequencing data, which we have shown to be a strong signal for CNA probabilities and selection parameters.

In contrast, the phylogeny tip statistics differ significantly between MEDICC2 and ground truth. These statistics require accurately segregating individual tips from the remaining of the sample. For instance, locating the two tips in a cherry requires at least one CNA that differentiates them from the other cells. A pitchfork likewise requires one CNA to distinguish a group of three tips. Similarly, identifying a ladder depends on a sequence of tips that are uniquely related to each internal node. Even for high levels of *p*_*misseg*_, the number of mis-segregations observed in a sample is typically not high enough to separate distinct phylogeny tips, resulting in the discrepency between MEDICC2 results and the true values. We note that there is evidence for cell-specific CNAs in recent scDNA data [6, 9, 10]. Most of these events are focal amplifications and deletions, and only a few are large events such as whole-chromosome or chromosome-arm missegregations. The cell-specific events can help identifying single tips in the phylogeny, refining the phylogeny tip statistics. Another potential approach is to utilize unique mutations in single cells. Because mutations occur at a much higher frequency than CNAs, they can be used to differentiate among distinct cells. However, due to low coverage, it is challenging to reliably detect unique mutations in scDNA-seq data. Future improvements in sequencing technologies and developements of phylogeny inference from both CNA and mutational data could increase the accuracy of phylogeny tip statistics.

Finally, we analyze the MEDICC2-based phylogeny balance statistics. Compared to the true values, the balance indices inferred from MEDICC2 phylogeny are mostly accurate, with the exception of the stairs index. In our correlation study, the stairs index has the weakest correlation with *p*_*misseg*_ among balance indices (Figure 3). The ABC-rf variable importance analysis also finds it to be a limited indicator for the missegregation probability (Figure S1).

In short, we find that all statistics based on CN profiles and most phylogeny balance indices in our study can be estimated reliably from MEDICC2, across different values of missegregation probabilities. In previous sections, we also found that these statistics are valuable in inferring CNA probabilities and selection parameters. On the other hand, phylogeny tip statistics cannot be reliably estimated from CN profiles, and do not carry a strong signal for the inference problem either.

## 3 Conclusion

In order to fully characterize the genomic evolution from DNA-seq data, it is important to quantify the rates at which CNAs arise and the selection forces that act upon them. Valind et al. constructed a discrete in silico model to analyze the increased prevalence of aneuploidy in cancer [43]. Their model assumes strong negative selection on aneuploid cells in normal tissues that is relaxed in tumors. While CNAs are indeed more tolerated in cancer, sequencing data has shown elevated frequencies of certain aneuploidies, either across tumors or specifically to some cancer subtypes [23], indicating that they are under positive selection. Elizalse et al. adapted a Markov-chain model to estimate the karyotypic distributions [44], building upon a stochastic chromosome copy number evolution model [45]. This approach derived optimal chromosome missegregation probabilities for maximal karyotypic heterogeneity while optimizing the computational efficiency. Salehi et al. employed a Bayesian fitness model grounded in the Wright-Fisher diffusion to infer clonal fitness coefficients from their growth trajectories in time-series scDNA-seq data [9]. Since the model does not consider new CNAs, it is applicable for the analysis of short-term dynamics and might be inefficient for studying the entire tumor history. Lynch et al. developed a model to infer both selection and CNA rates from scDNA-seq [19]. The framework models fitness as a scaling factor multiplied by the OG-TSG score from Davoli et al. [11], which quantifies the count and potency of oncogenes and tumor suppressor genes, then infers the scaling factor and missegregation rate. As the OG-TSG score is computed for all genes across pan-cancer TCGA samples, it might be challenging to adapt this framework to study individual cancer types, where identification of driver genes and their mutation frequencies is difficult due to low sample counts.

We have recently introduced CINner [18], a simulation framework that models the impact of CNA probabilities and tissue-specific selection coefficients on the observed karyotypes. In this paper, we investigate the problem of inferring these parameters, which together shape the aneuploidy patterns in cancers. Cancers with high CNA probabilities are expected to be more heterogeneous [18], which has been linked to poor patient outcome and higher chance of relapse [46] Therefore, accurate parameter inference promises to aid in stratifying patient subtype and predict tumor progression. However, reliable estimation of both CNA rates and selection parameters from data is challenging. We have previously shown that these variables impact most bulk DNA-seq statistics simultaneously, making it difficult to correctly infer each [18]. For instance, a frequently observed CN gain could be either because the chromosome harbors many oncogenes (selection parameter ≫ 1) or because missegregations regularly occur in the cancer type (*p*_*misseg*_ ≫ 0). Likewise, high tumor heterogeneity could be due to high CAN probabilities or most chromosomes acting neutrally (selection parameter ≈ 1), which results in many subclones co-existing in the absence of selection.

To solve the nonidentifiability issue, we construct an inference method that takes as input a mixture of bulk and single-cell DNA-sequencing data. The algorithm is based on Approximate Bayesian Computation [26, 29] and CINner [18]. We consider several statistics based on CN profiles and cell phylogeny, which can be readily estimated from genomic data with current technologies. We find that statistics depicting clonal heterogeneity and CNA levels in sample CN profiles, together with indices for the degree of balance in scDNA-seq cell phylogenies, are important in estimating CNA probabilities. In contrast, only statistics for CN profiles are effective for inferring selection parameters. This is likely due to the fact that one chromosome’s selection parameter has negligible impact on the whole sample phylogeny. In both cases, there is little signal in the statistics quantifying local features of the phylogeny (e.g. cherries, pitchforks, and ladders).

We show that our algorithm can accurately recover both missegregation probability and chromosome-specific selection parameters from DNA-seq data. Peak performance requires at least 50 bulk DNA samples, which already exists for many cancer types [2, 1]. However, most available scDNA studies contain less than the minimum of 20 or 40 samples necessary to minimize the uncertainty and error in the inference, respectively [6, 9, 10]. Nevertheless, the higher errors associated with employing smaller scDNA-seq cohorts still appear adequately modest, therefore biologically meaningful interpretations can still be gained. Furthermore, our numerical experiment with MEDICC2 shows that the statistics most important in inferring missegregation probability and the selection forces can be estimated accurately from real DNA-seq data.

In this study, we consider only whole-chromosome missegregations. However, it is straight-forward to expand the framework to consider other CNA mechanisms simultaneously, such as chromosome-arm missegregations and whole-genome duplication [47, 14]. In such cases, more detailed copy number segmentation and phylogeny reconstruction are necessary to distinguish between different CNA events. CINner [18] is a prominent tool for this purpose, since additional chromosomal instability mechanisms can be incorporated with relative ease. The combination of CINner and the inference framework described in this paper provides a promising approach in analyzing existing and upcoming cancer datasets, as both bulk and single-cell DNA-sequencing become more readily available.

## Code availability

All code for parameter inference, data analysis and sensitivity studies is available at https://github.com/dinhngockhanh/CINner_missegregation_inference.

## Acknowledgments

The authors acknowledge the support from the Herbert and Florence Irving Institute for Cancer Dynamics and Department of Statistics at Columbia University.

## Additional information

The authors declare no competing interests.

## Author contributions

K.D. conceived the presented concept and methodology; Z.X., Z.L. and K.D. wrote the code; Z.X. and Z.L. carried out the analysis and wrote the original manuscript draft; K.D. supervised, reviewed and edited the final manuscript draft.

## Methods

### Mathematical model of CINner

We simulate the clonal evolution with CINner [18]. For convenience, in this section, we summarize the mathematical model and parameters of interest.

For this study, CINner is restricted to copy numbers of whole autosomes. Each chromosome *i* = 1, …, 22 has selection parameter *s*_*i*_ ∈ (0, ∞), which is constant among different cells. The CN profile of each cell *k* can be characterized as a vector 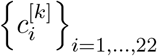, where 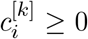 is the count of chromosome *i*. The cell’s fitness is then defined as

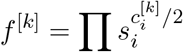

However, if 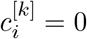 for any chromosome *i* (that is, the cell contains nullisomy), then *f* ^[*k*]^ = 0. Each simulation starts with *N* (0) diploid cells, i.e.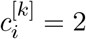. Each cell follows a birth-death process. Its lifespan is exponentially distributed with the same rate *λ*. At the end of its life, the cell either divides or dies. Suppose the population at time *t* consists of *P* (*t*) cells. The probability that cell *k* divides is

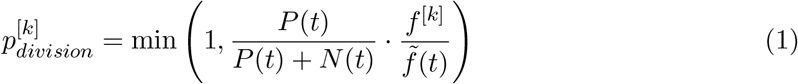

where 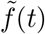 is the average fitness among cells present at time *t*:

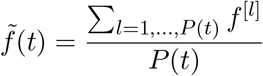

Note that (1) ascertains that on average, the total cell population *P* (*t*) follows a pre-determined dynamic *N* (*t*).

If cell *k* divides, a missegregation occurs with probability *p*_*misseg*_. If a missegregation event happens, a uniformly random chromosome *j* is chosen. The CN profiles 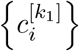 and 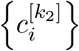 of the two progeny cells *k*_1_ and *k*_2_ are then defined as:

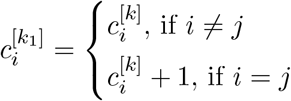

and

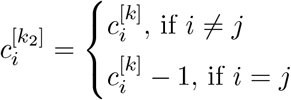

that is, cell *k*_1_ gains another copy of chromosome *j* from parent cell *k*, and cell *k*_2_ loses that copy.

The model therefore has two fixed variables:

- Mean cell lifespan *λ* = 30 days 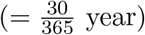.
- Expected total population dynamic 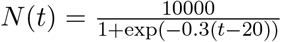, which is a logistic function plateauing to 10,000 cells with the midpoint at 20 years. Simulated bulk or scDNA-seq samples consist of 1,000 cells randomly chosen at *t* = 80 years. and two parameters which are targets for inference:
- Probability of missegregation *p*_*misseg*_
- Chromosome selection parameters {*s*_*i*_}

### Statistics for bulk and single-cell DNA sequencing data

#### Bulk CN distance

The bulk DNA-seq data cohort consists of *N*_*bulk*_ samples. Each sample *k* = 1, …, *N*_*bulk*_ is associated with a CN profile 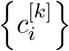.

For each parameter set *θ* = {*p*_*misseg*_, *s*_1_, …, *s*_22_}, *N*_*bulk*_ simulations are created with CINner, resulting in *N*_*bulk*_ CN profiles, where each sample *l* is associated with CN profile 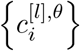. We seek to define a distance between 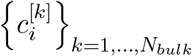 and 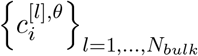, using the Wasser-stein distance [48, 49]. Shortly, Wasserstein(*µ* → ν; *C*) measures the minimum cost of moving mass from probability distribution *µ* to probability distribution ν, given a cost function *C* such that *C*(*x, y*) is the cost of transporting a unit from point *x* in *µ* to point *y* in ν.

We define the cost matrix 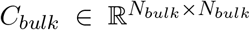 such that each entry is the Euclidean distance between corresponding data sample and simulated sample:

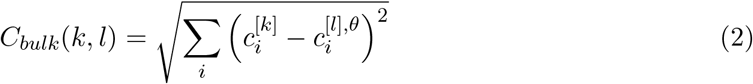

We then compute the CN distance between 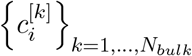 and 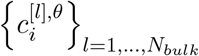 by using function `wasserstein` in `R` library `transport` with power `p`=1 [32] to find the Wasserstein cost of transforming the data sample distribution 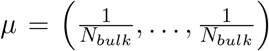 into the simulated sample distribution 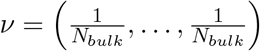, subject to cost matrix *C*_*bulk*_ (Figure 2b):

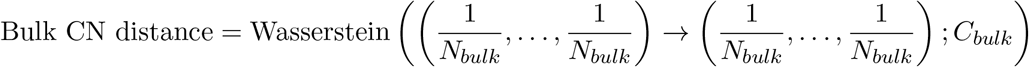

In this study, we simulate the bulk DNA-seq data cohort also with CINner. That is, we choose ground-truth parameters *θ*^[*truth*]^ and simulate 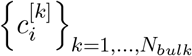. We expect that the bulk CN distance decreases as *θ* → *θ*^[*truth*]^, as the simulated samples have similar gains and losses on the same chromosomes as in the data.

The choice of metric between two samples could potentially affect the performance of the bulk CN distance in capturing the signals from data. We choose Euclidean distance (2) because it is a metric, hence the Wasserstein distance is also a metric. Other more biologically relevant distances such as [21, 50, 41], which are based on the minimal number of CNAs required to transform one CN profile into another, do not always satisfy metric considerations (e.g. *C*_*bulk*_(*k, l*)≠ *C*_*bulk*_(*l, k*)), therefore their applications should be taken with consideration.

#### Single-cell CN distance

The single-cell DNA-seq data cohort consists of *N*_*sc*_ samples. Each sample *m* = 1, …, *N*_*sc*_ contains *N*_*m*_ cells, and each cell *k* = 1, …, *N*_*m*_ is associated with a CN profile 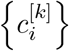.

We create *N*_*sc*_ simulations using CINner with a given parameter set *θ*. Each sample *n* has *N*_*n*_ cells, where each cell *l* has CN profile 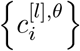. Similarly to the bulk CN distance, the single-cell CN distance is defined as the Wasserstein cost of transforming the data sample distribution 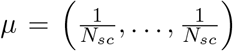 into the simulated sample distribution 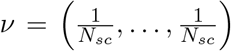, subject to cost matrix 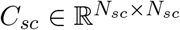 (Figure 2c):

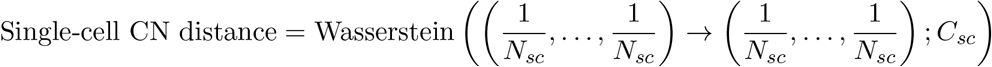

Distance *C*_*sc*_(*m, n*) between data sample *m* and simulated sample *n* is defined as the Wasserstein distance between the single-cell CN profiles from the data sample 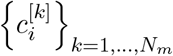 and the simulated sample 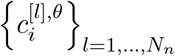:

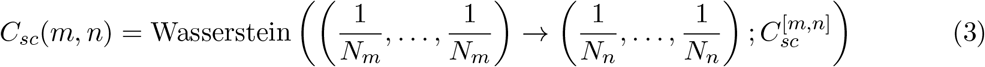

where 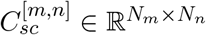 stores the CN distance between each pair of cells:

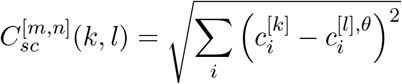

In total, for each parameter set *θ*, this procedure requires 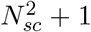 runs of the `wasserstein` function. Another consideration is that recent single-cell DNA samples contain up to thousands of cells, rendering (3) extremely computationally expensive. To optimize the runtime, we take advantage of the fact that the cells in the sample can often have similar if not identical CN profiles. Assume that cells in data sample *m* can be grouped into *C*_*m*_ clones, such that all *N*_*α*_ cells in clone *α* have the same CN profile 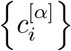 and different clones have different CN profiles 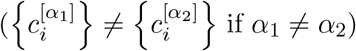. Similarly, simulated sample *n* can be clustered into *C*_*n*_ clones, where clone *β* consists of *Ñ*_*β*_ cells of CN profile 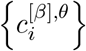. The distance between samples *m* and *n* (3) can then be replaced with:

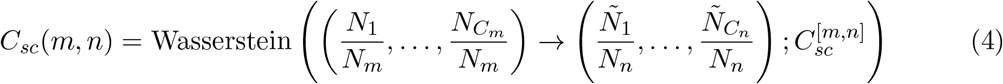

where 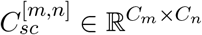 contains the CN distances between each pair of clones:

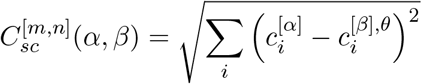

Because the clone count is often much smaller than cell count, i.e. *C*_*m*_ ≪ *N*_*m*_ and *C*_*n*_ ≪ *N*_*n*_, (4) results in the same value as (3) with significantly reduced runtime.

Similar to the bulk DNA data, in this study, we simulate the single-cell CN data assuming ground-truth parameters *θ*^[*truth*]^. We consider *N*_*m*_ = *N*_*n*_ =1,000 cells in each data and simulated sample.

#### Shannon diversity index

Given a single-cell DNA-seq sample *m*, clustered into *C*_*m*_ distinct clones where there are *N*_*α*_ cells of each clone *α*. Its Shannon diversity index is defined as

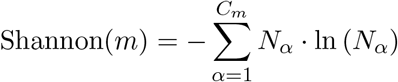

computed with function `diversity` in R library `vegan` [51]. We consider two statistics for the whole scDNA-seq cohort consisting of *N*_*sc*_ samples: mean and variance of Shannon(m) across all samples. These values are compared against the mean and variance of Shannon indices from CINner samples with varying *θ* in our ABC-based parameter inference, to be described below.

For the remaining statistics, we likewise use their mean and variance across the corresponding DNA-seq cohort in this study. Therefore, in the following sections, we will describe only how to derive the statistic from one DNA-seq sample.

#### Missegregation count in bulk DNA-seq samples

We assume that the CN profile in each bulk DNA-seq sample represents the average CN profile among all sampled cells. Therefore, we define Missegregation count(k) for sample k to be the average number of missegregations that a sampled cell has acquired since *t* = 0.

#### Clonal and subclonal missegregation counts in scDNA-seq samples

Given a scDNA-seq sample *m*, let *𝒯*_*m*_ be the phylogeny tree with *N*_*m*_ leaves, each of which corresponds to a cell in the sample. The tree traces the merging of these cells back to a most recent common ancestor (MRCA). Any observed missegregation event *γ* can be placed on an edge in *𝒯*_*m*_, let *N*_*γ*_ be the number of cells descending from this edge (0 < *N*_*γ*_ ≤ *N*_*m*_).

The clonal missegregation count (Figure 2a) is defined to be the number of missegregations that the MRCA has acquired. Alternatively, any clonal event must be present in all cells, therefore the clonal missegregation count is the number of events *γ* such that *N*_*γ*_ = *N*_*m*_:

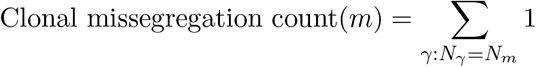

On the other hand, a subclonal event is carried by only a subset of sampled cells. The subclonal missegregation count (Figure 2a) is weighted by number of cells carrying each event:

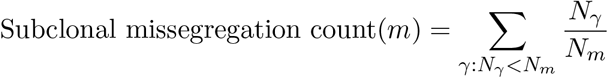

#### Cherry count, pitchfork count, IL number and average ladder of cell phylogeny

We now define the phylogeny tip statistics for the scDNA-seq tree *𝒯*_*m*_ with *N*_*m*_ leaves (Figure 2a). The computations are performed with R library `PhyloTop` [34]. For rooted binary trees, a cherry is a pair of leaves that are adjacent to a common ancestor node [52]. The cherry count is the normalized number of cherries *c*(*𝒯*_*m*_) [35], computed with function `cherries`:

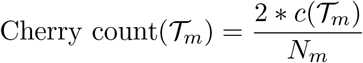

A pitchfork is a subtree with three leaves [35]. The pitchfork count is the normalized number of pitchforks *p*(*𝒯*_*m*_), computed with function `pitchforks`:

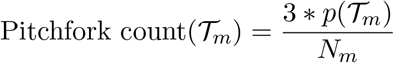

An IL node is an internal node in *𝒯*_*m*_ that has a single leaf child [35]. The IL number is the count of IL nodes normalized for leaf count, computed with function `ILnumber`:

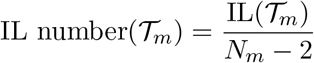

A ladder is a path composed only of IL nodes. The average ladder is the average ladder length AL(*𝒯*_*m*_) normalized for leaf count, computed with function `AvgLadder`:

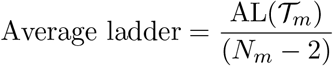

#### Balance indices of cell phylogeny

Balance indices measure the degree of balance or imbalance in the phylogeny tree *𝒯*_*m*_ with *N*_*m*_ leaves (Figure 2a). Stairs calculates the proportion of subtrees where two child branches have different number of descendants [53], computed with function `stairs` in R library `PhyloTop` [34].

Colless index is the sum of absolute differences between number of leaves descending from the left and right branches of each internal node in *𝒯*_*m*_, normalized by dividing by 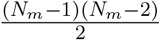 [54]. We compute the Colless index with function `colless.phylo` in R library `PhyloTop` [34].

Sackin index is the sum of ancestor counts from all leaves to the MRCA [55, 35], normalized by dividing by 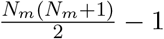 [54] (function `sackin.phylo` in R library `PhyloTop` [34]).

B2 index is computed with function `B2I` in R library `treebalance` [35], and measures the probability of reaching the leaves in *𝒯*_*m*_ when starting at the MRCA and taking a random branch at each internal node with equal probabilities:

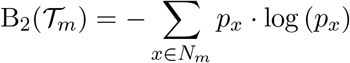

where *p*_*x*_ is the probability of ending at leaf *x* with ancestor set ancestor(*x*), each node *v* in which has child(*v*) childen:

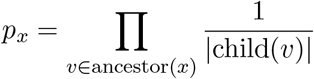

Finally, Max depth is the maximal depth for all vertices in *𝒯*_*m*_, computed with function `maxDepth` in R library `treebalance` [35].

### Parameter inference with Approximate Bayesian Computation

#### Simulated data

We simulate the DNA-seq data that is used to infer back the missegregation probability and chromosome selection parameters. The ground truth parameters are:

- *p*_*misseg*_ = 2 × 10^−4^
- *s*_*i*_ ∼ Uniform, 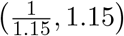

We then use CINner to simulate *N*_*bulk*_ = 100 bulk DNA-seq samples and *N*_*sc*_ = 50 scDNA-seq samples. Each simulation consists of 1,000 cells sampled at time *t* = 80 years. Finally, we compute *𝒮*^[𝒟]^, consisting of statistics from the bulk and single-cell data cohort as described previously.

#### ABC library

We prepare a simulation library to train ABC-rf (to be discussed next) to infer model parameters from the data. The library consists of 100,000 simulations. For each simulation, we perform the following:

1. Sample log_10_ (*p*_*misseg*_) ∼ Uniform(−5, −3). Sample *s*_*i*_ ∼ Uniform 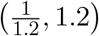.
2. Create *N*_*bulk*_ + *N*_*sc*_ CINner simulations with chosen parameters.
3. Compute statistics.

The library then consists of 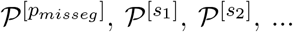 and *𝒮*^[ℒ]^, such that the *j*th simulation assumed parameter set 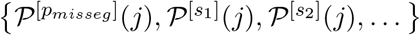 and resulted in statistics *𝒮*^[ℒ]^(*j, ·*).

We group the statistics into three categories:

- CN statistics: Bulk CN distance, single-cell CN distance, mean and variance of Shannon diversity index, missegregation count in bulk DNA-seq samples, clonal and subclonal missegregation count in scDNA-seq samples.
- Phylogeny tip statistics: mean and variance of cherry count, pitchfork count, IL number, average ladder.
- Phylogeny balance statistics: mean and variance of stairs, Colless index, Sackin index, B2 index, max depth.

Depending on the chosen categories, corresponding statistics from the simulations are compiled into the ABC library *𝒮*^[ℒ]^.

The CN statistics are computed in two distinct ways. In the first method, the statistics are measured using the entire genome-wide CN vectors from each data and simulated sample. In the second method, the CN information is restricted to a given chromosome *i*.

#### Parameter inference with ABC-rf

We apply ABC random forect (ABC-rf, [29]) to infer model parameters from the data and training library. For each parameter *θ* (= *p*_*misseg*_, *s*_1_, *s*_2_, …), we do the following:

1. Train the random forest: use function `regABC-rf` to train model *ℳ* on 𝒫 ^[*θ*]^ ∼ *𝒮*^[ℒ]^.
2. Infer the posterior distribution *π*(*θ*): by applying function `predict` on model *ℳ* and data statistics *𝒮*^[𝒟]^.

For *p*_*misseg*_, the CN statistics in *𝒮*^[ℒ]^ and *𝒮*^[𝒟]^ are based on genome-wide CN vectors, described above. For selection parameters *s*_*i*_, the CN statistics based on chromosome *i* are used. Figure S1 presents the ABC-rf evaluations of the trained random forest for the missegregation probability *p*_*misseg*_. Figures S2 and S3 show the evaluations for *s*_1_ and *s*_2_, the selection parameters of chromosomes 1 and 2.

**Figure S1:**
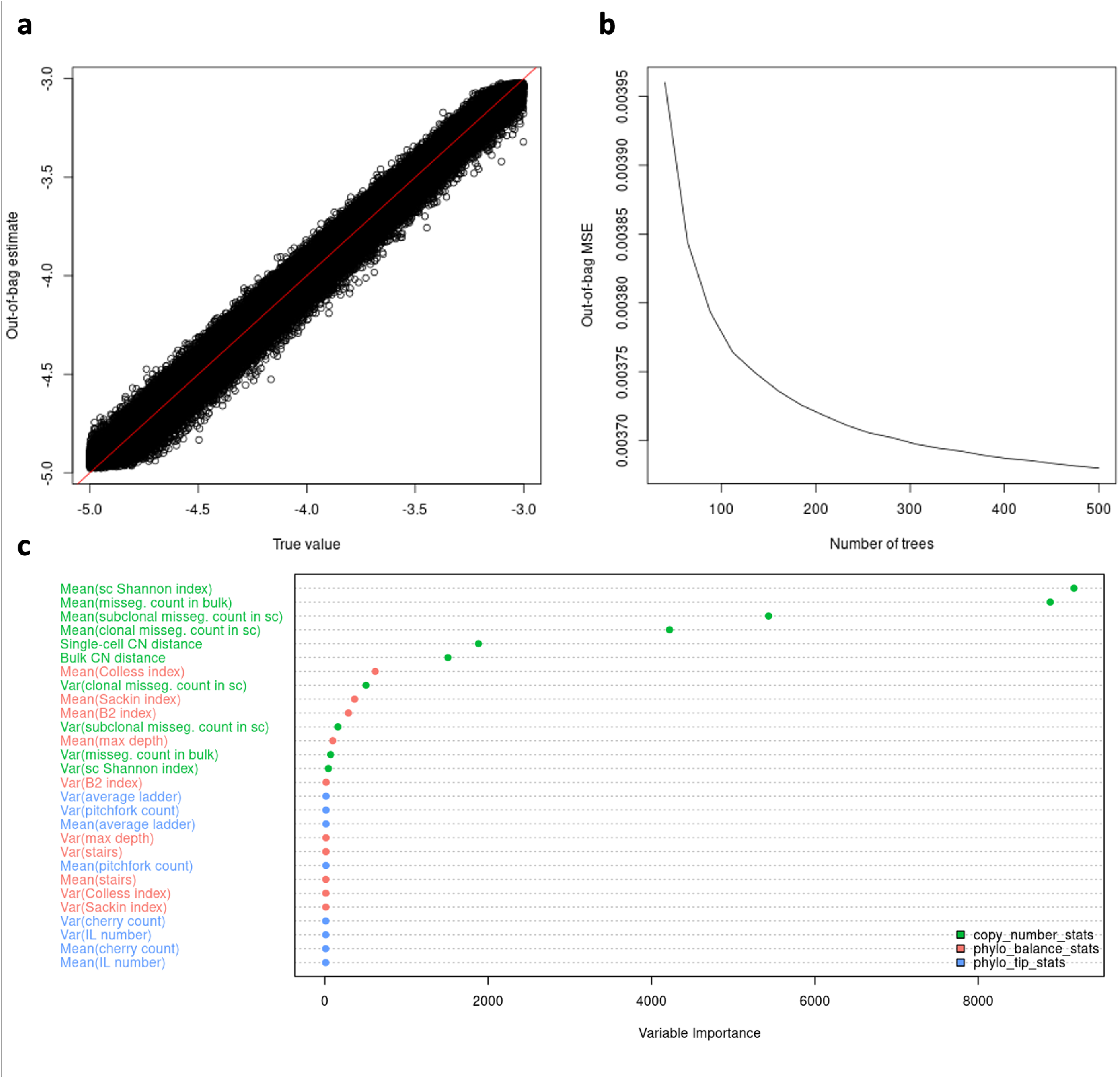
ABC-rf reports for the inference of *p*_*misseg*_. **a**: Out-of-bag estimates against true values of *p*_*misseg*_ in the training set. The red diagonal line indicates where the estimate is identical to the true value. **b**: Relationship between out-of-bag Mean Squared Error (MSE) and number of trees in the ABC-rf random forest. **c**: Analysis of variable importance among statistics employed in ABC-rf to infer *p*_*misseg*_. Statistics are categorized into CN statistics (green), phylogeny tip statistics (blue) and phylogeny balance statistics (red).

**Figure S2:**
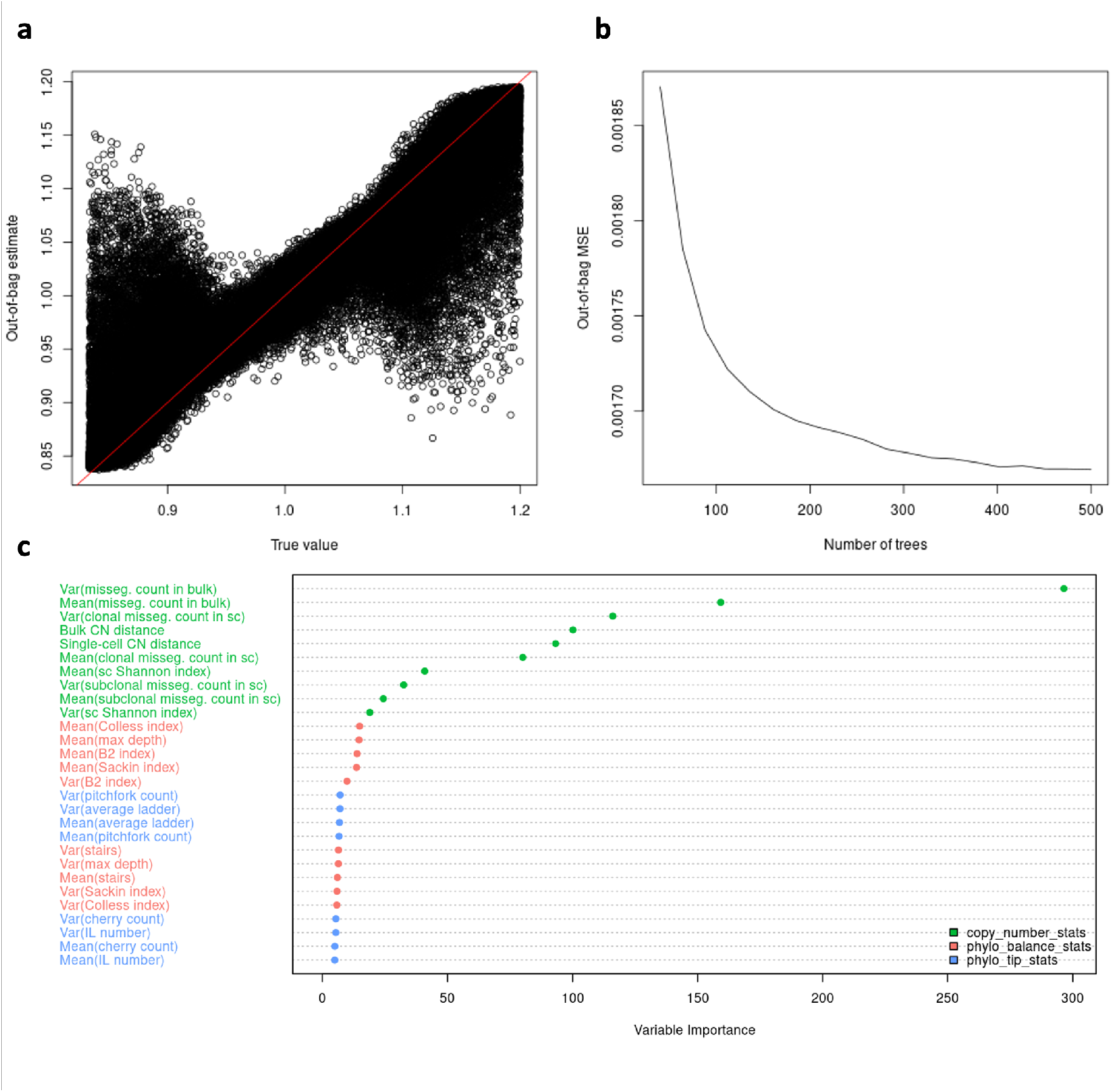
ABC-rf reports for the inference of *s*_1_. **a**: Out-of-bag estimates against true values of *s*_1_ in the training set. The red diagonal line indicates where the estimate is identical to the true value. **b**: Relationship between out-of-bag Mean Squared Error (MSE) and number of trees in the ABC-rf random forest. **c**: Analysis of variable importance among statistics employed in ABC-rf to infer *s*_1_. Statistics are categorized into CN statistics (green), phylogeny tip statistics (blue) and phylogeny balance statistics (red).

**Figure S3:**
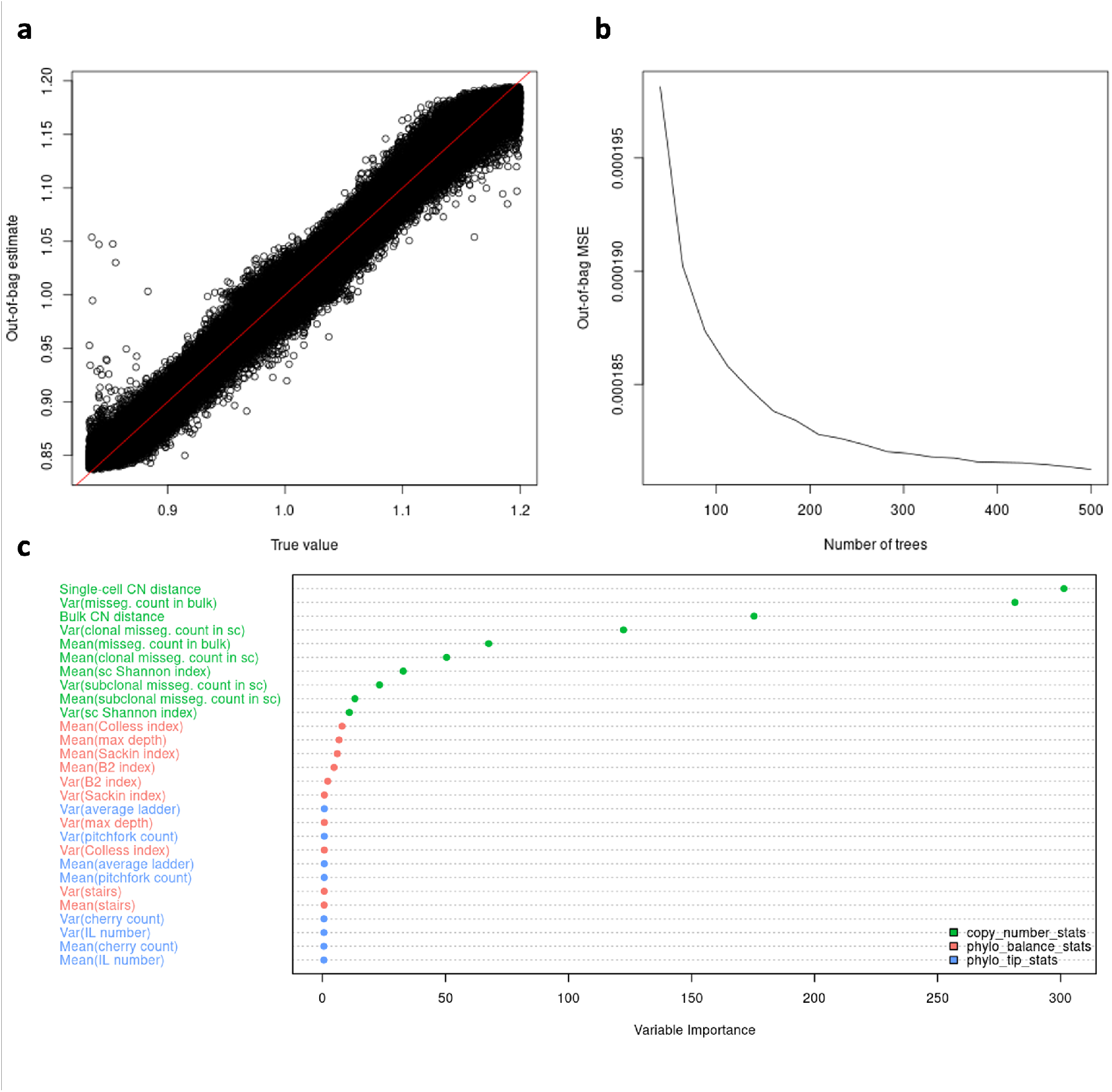
ABC-rf reports for the inference of *s*_2_. **a**: Out-of-bag estimates against true values of *s*_2_ in the training set. The red diagonal line indicates where the estimate is identical to the true value. **b**: Relationship between out-of-bag Mean Squared Error (MSE) and number of trees in the ABC-rf random forest. **c**: Analysis of variable importance among statistics employed in ABC-rf to infer *s*_2_. Statistics are categorized into CN statistics (green), phylogeny tip statistics (blue) and phylogeny balance statistics (red).

#### Analysis of inference

We characterize the accuracy and certainty of the inference in several ways. For each parameter *θ*, we find the mean, median, mode, and standard deviation of the posterior distribution *π*(*θ*).

To summarize the inference for all chromosome selection parameters, we compute the average standard deviation among *π*(*s*_*i*_) for *i* = 1, …, 22, and the Root Mean Square Error (RMSE):

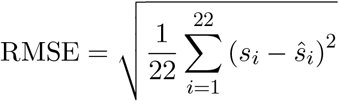

where *s*_*i*_ and *ŝ*_*i*_ are the true value and posterior mean of selection parameter of chromosome *i*, respectively.

### Correlations between statistics and model parameters

For a given combination of parameter *θ* and statistic *s*, we compute the Pearson correlation from the 100,000 simulations in the ABC library by using function cor in R. Similar to the ABC inference, for CN statistics, the genome-wide values are used to compute the correlations against *p*_*misseg*_, and the values computed for chromosome *i* are compared against each *s*_*i*_.

### Sensitivity analysis with respect to sample sizes

To study the sensitivity of the inference method with respect to the bulk DNA-seq sample size, we keep *N*_*sc*_ = 50 and vary *N*_*bulk*_ ∈ {10, 20, …, 100}. We then restrict the sample size when computing *𝒮*^[𝒟]^ and *𝒮*^[ℒ]^ according to the value of *N*_*bulk*_.

Similarly, we fix *N*_*bulk*_ = 100 and vary *N*_*sc*_ ∈ {5, 10, …, 50} to find the number of scDNA-seq samples necessary to minimize both RMSE and standard deviation in the ABC posterior distributions.

### Accuracy of statistics inferred from MEDICC2

For each simulation, we first sample log (*p*_*misseg*_) from Uniform (5, 3) and chromosome selection parameters *s*_*i*_ from Uniform 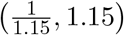. One CINner simulation is then created, resulting in the CN profiles for each sampled cell and the true phylogeny tree. We then use MEDICC2 [21, 50] to infer the phylogeny from single-cell CN profiles. The statistics computed from the MEDICC2 phylogeny are then compared against statistics from the true phylogeny.

We also directly compare the true and MEDICC2 phylogenies, using the generalized Robinson-Foulds distance (function `TreeDistance` in `R` library `TreeDist` [56]).

## Notes

https://github.com/dinhngockhanh/CINner_missegregation_inference

### Competing Interest Statement

The authors have declared no competing interest.

### Summary of Updates

We changed one sentence in the Abstract that might have sounded ambiguous.

